# SV2A-Syt1 interaction controls surface nanoclustering and access to recycling synaptic vesicles

**DOI:** 10.1101/2021.12.08.471864

**Authors:** Christopher Small, Callista Harper, Christiana Kontaxi, Elizabeth Davenport, Tristan Wallis, Anusha Malapaka, Nyakuoy Yak, Merja Joensuu, Ramon Martínez-Mármol, Michael A. Cousin, Frédéric A. Meunier

## Abstract

Following exocytosis, the recapture of vesicular proteins stranded at the plasma membrane in recycling synaptic vesicles (SVs) is essential to sustain neurotransmission. Nanoclustering is emerging as a mechanism through which proteins may be ‘pre-assembled’ prior to endocytosis, to ensure high fidelity of retrieval for subsequent rounds of vesicle fusion. Here, we used single molecule imaging to examine the nanoclustering of synaptotagmin-1 (Syt1) and synaptic vesicle protein 2A (SV2A). Syt1 forms surface nanoclusters through interaction of its C2B domain (K326/K328) with SV2A, as demonstrated by mutating Syt1 (K326A/K328A) and knocking down endogenous SV2A. Blocking cognate interaction with Syt1 (SV2A^T84A^) also decreased SV2A clustering. Impaired nanoclustering of Syt1 and SV2A leads to accelerated endocytosis of Syt1, altered intracellular sorting and decreased trafficking of Syt1 to a Rab5-positive endocytic pathway. We conclude that the interaction between SV2A and Syt1 locks both molecules into surface nanoclusters, controlling their entry into recycling SVs.

## Introduction

Synaptic vesicle (SV) recycling involves a balance between fusion (exocytosis) and retrieval (endocytosis) of SVs from the plasma membrane (PM) at nerve terminals during neurotransmission. Both exocytosis and compensatory endocytosis involve the coordinated actions of proteins and lipids to ensure high fidelity at high rates of fusion. To sustain neurotransmission, recycling SVs need to recapture essential vesicular machinery stranded at the PM. However, the mechanisms through which neurons retrieve essential vesicular proteins from the PM are not well defined. As certain SV proteins lack canonical recognition motifs for endocytic adaptor molecules, interactions between vesicular cargoes may facilitate recruitment of proteins from the PM into SVs (Gordon and Cousin, 2016), preserving vesicle protein stoichiometry (Takamori et al., 2006; Wilhelm et al., 2014) during neurotransmission. For example, vesicle-associated membrane protein 2 (VAMP2) is a soluble N-ethylmaleimide-sensitive factor attachment protein receptor (SNARE) that regulates fusion of SVs with the PM (Südhof and Rothman, 2009) and its internalisation is facilitated in part, via its interaction with synaptophysin (Gordon and Cousin, 2013; Gordon et al., 2011; Harper et al., 2021; Harper et al., 2017). Similarly, vesicular glutamate transporter 1 (vGlut1) facilitates the recruitment of multiple SV proteins from the PM back into SVs (Pan et al., 2015). Thus, interactions between vesicular molecules are theorised to improve the fidelity of endocytic uptake and allow SVs to retain their protein organization during multiple rounds of fusion.

One mechanism through which protein interactions improve the fidelity of endocytosis is nanoclustering. Following exocytosis, vesicular proteins stranded on the PM cluster through protein-protein interactions: notably, VAMP2 disperses following exocytosis and subsequently re-clusters via interactions with endocytic proteins, particularly AP180 and CALM (Gimber et al., 2015). The endocytic machinery therefore has the potential to initiate clustering of surface stranded vesicular proteins. However, it is not clear what factors control clustering of other vesicular proteins, such as synaptotagmin-1 (Syt1). Syt1 is a transmembrane SV molecule involved in calcium (Ca^2+^)-dependent exocytosis (Geppert et al., 1994) and clusters at the PM (Opazo et al., 2010; Willig et al., 2006). Syt1 binds to phosphoinositol(4,5)bisphosphate in a Ca^2+^-dependent manner through its cytosolic C2A and C2B domains (Bai et al., 2002; Schiavo et al., 1996; Stein et al., 2007) to mediate exocytosis. Syt1 forms a complex with another vesicular transmembrane protein, synaptic vesicle protein 2A (SV2A) (Bennett et al., 1992), which comprises twelve transmembrane-spanning domains capped by cytosolic C-terminal and N-terminal regions. The SV2A-Syt1 interaction occurs via the cytosolic domains of Syt1 and SV2A: hydrogen bonds are formed between two lysine residues (K326/K328) residing in the polybasic region of Syt1’s calcium-binding C2B domain (Fernandez et al., 2001) and the T84 epitope on the N-terminus of SV2A upon phosphorylation of the T84 epitope by casein kinase I (Zhang et al., 2015). SV2A has an enigmatic function in neurotransmission, which involves controlling the trafficking of Syt1 in neurons. Notably, the absence of SV2A reduces Syt1 levels in SVs and at the PM (Yao et al., 2010). SV2A interacts with Syt1 following membrane fusion (Wittig et al., 2021) and controls the retrieval of Syt1 during endocytosis (Kaempf et al., 2015; Zhang et al., 2015). For these reasons, SV2A is also a strong candidate as a regulator of Syt1 nanoclustering during SV recycling.

In this study, we investigated the role of protein-protein interactions in controlling the clustering of vesicular machinery at the PM, and how protein interaction and clustering events facilitate entry of proteins into recycling SVs during endocytosis. We hypothesised that nanoclustering of vesicular proteins at the PM allows for the generation of a ‘readily-accessible’ pool of pre-assembled molecules, forming a depot from which to selectively retrieve vesicular proteins into nascent recycling SVs. Using super-resolution imaging, we identified the determinants of Syt1 and SV2A nanoclustering by manipulating interactions between Syt1 and SV2A, as well as interactions with endocytic machinery. The Syt1-SV2A interaction was critical for their respective surface nanoclustering, with manipulation of the endocytic machinery having no effect on the nanoclustering of either molecule. Blocking SV2A-Syt1 nanoclustering accelerated Syt1 retrieval during SV endocytosis. This manipulation also led to increased mobility of internalised Syt1 suggesting alterations in SV nanoscale organization. The findings presented in this study suggest that Syt1 is dynamically sequestered into nanoclusters in an activity-dependent manner through its interaction with the N-terminal tail of SV2A, and that the nanoclustering of Syt1 by SV2A decreases the kinetics of Syt1 endocytic uptake. Accordingly, we report that SV2A interaction also controls the organization of Syt1 following internalisation, causing Syt1 entrapment within endocytic pathways associated with early endosome formation. SV2A therefore plays a critical role in the recycling of Syt1, with implications for the function of Syt1 during multiple rounds of vesicle fusion.

## Results

### The Syt1 C2B domain (K326/K328) interaction with SV2A controls the activity-dependent confinement of plasma membrane-stranded Syt1

First, we investigated the surface mobility and nanoclustering of Syt1 in primary cultures of hippocampal neurons using universal Point Accumulation Imaging in Nanoscale Topography (uPAINT) imaging. This single-particle tracking technique allows selective analysis of the nanoscale organisation of surface proteins via labelled ligand tracking (Giannone et al., 2010; Giannone et al., 2013; Joensuu et al., 2016). We overexpressed Syt1 tagged with pHluorin (Syt1-pH), a pH-sensitive green fluorescent protein (GFP) in primary cultures of mouse hippocampal neurons. The pHluorin (pH) moiety is quenched in the acidic SV environment and unquenched following exocytosis due to exposure to the neutral, extracellular pH (Miesenböck et al., 1998). The epifluorescence of Syt1-pH revealed distinct synaptic boutons lining the axon of mature neurons (Fig. 1A i-ii). To track single molecules of Syt1-pH on the PM, we applied atto647N-labelled anti-GFP nanobodies (NBs) (Gormal et al., 2020; Kubala et al., 2010) in a depolarizing buffer (56 mM K^+^) to increase SV recycling and Syt1-pH PM levels. The atto647N fluorophore was excited (647 nm) in total internal reflection fluorescence (TIRF) (50 Hz), to selectively image the surface population of atto647N-NB-bound Syt1-pH (16000 frames, 320 s) (Fig. 1A iii-v). The effect of ablating SV2A interactions on the nanoscale organization of Syt1 was also examined. We expressed a mutant form of Syt1 (Syt1^K326A/K328A^-pH) (Fig. 1B i-ii) containing two lysine (K) to alanine (A) substitutions in the polybasic region of Syt1’s C2B domain that have been shown to block interaction with the cytosolic, N-terminus of SV2A (Borden et al., 2005). A loss of subsynaptic clustering of Syt1 was observed (Fig. 1B iii-v) in the presence of the K326A/K328A mutation (Fig. 1C i). As clustered Syt1 appeared prominently during stimulation, we imaged Syt1^WT^-pH-atto647NB before and after stimulation across the total hippocampal axon, to determine if clustering was activity-dependent. Under resting and stimulated conditions, the mean-square displacement (MSD) of Syt1-pH molecules was plotted over time (200 ms) (Fig. 1C ii). This was quantified by calculating the area under the MSD curve (AUC) and the ratio of mobile:immobile molecules (M/MM) (Fig. 1C iii). Both metrics were significantly decreased in response to stimulation (p=0.018 and p=0.008 respectively) demonstrating that confinement of Syt1 molecules on the PM was triggered in response to stimulation. However, in the presence of the K326A/K328A mutant which prevents SV2A binding to Syt1, the mobility of Syt1 remained constant, with no change in MSD AUC or M/MM (Fig. 1C iv). We concluded that the Syt1/SV2A interaction was critical for the activity-dependent confinement of Syt1 to surface nanoclusters.

**Figure 1.**
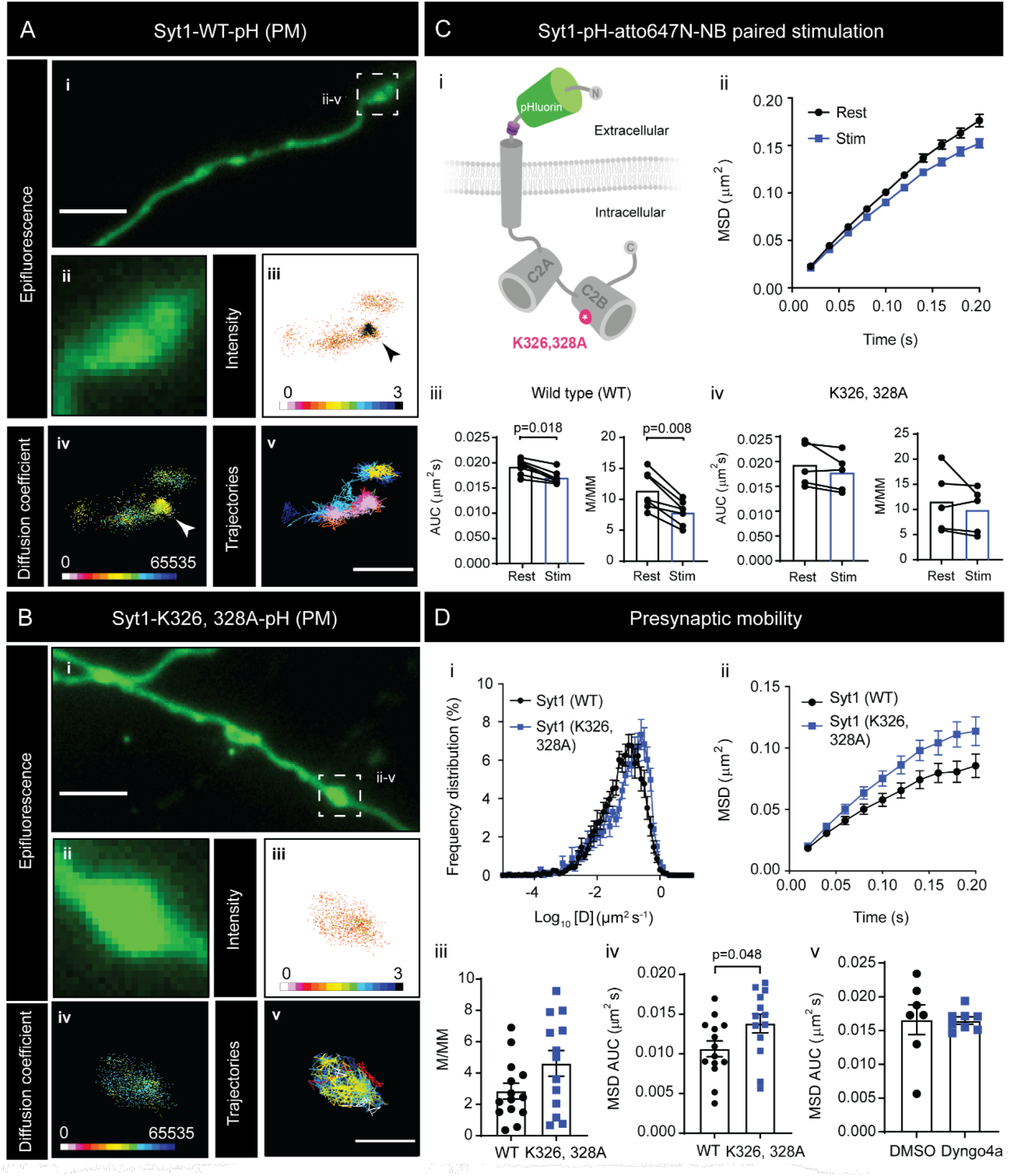
Inhibition of SV2A interaction (K326A/328A) increases the mobility of Syt1 following stimulation. Universal Point Accumulation for Imaging in Nanoscale Topography (uPAINT) of Syt1-pH at the plasma membrane (PM) was performed in hippocampal neurons treated with anti-GFP atto647N-nanobodies (NB). **(A)** Syt1^WT^-pH epifluorescence in (i) axons and (ii) nerve terminals. Super-resolved syt1-pH-atto647N-NB in nerve terminals highlighted by (iii) intensity, (iv) diffusion coefficient (D Coeff) and (v) trajectory maps. Arrows designate Syt1 hotspots. **(B)** Syt1^K326A/K328A^-pH epifluorescence in (i) axon (ii) nerve terminals. Superresolved Syt1^K326A/K328A^-pH-atto647-NB highlighted by (iii) intensity, (iv) D Coeff and (v) trajectories. Axon scale bar = 4 μm (A i, B i), presynapse scale bar = 1 μm (A v, B v). **(C)** (i) The K326A/K328A mutations (red) are located in the intracellular C2B domain of Syt1. The TEV peptide sequence (purple) and pH tag (green) are shown. (ii-iii) Syt1^WT^ imaged under resting and stimulated (high K^+^; 56 mM) conditions. Corresponding (ii) mean square displacement (MSD; μm^2^ over 200 ms), (iii, left) a significant decrease in the area under the curve (AUC) of MSD and (iii, right) M/MM. (iv) Syt1^K326A/328A^ under resting and stimulated (high K^+^; 56 mM) conditions. Corresponding (iv, left) MSD AUC (μm^2^s) and (iv, right) M/MM. **(D)** Presynaptic mobility of Syt1^WT^ and Syt1^K326A/328A^ shown by (i) Log_10_ D Coeff [D] (μm^2^s^-1^) frequency distribution (ii) MSD (μm^2^ over 200 ms), (iii) M/MM, (iv) MSD AUC (μm^2^s) and (v) MSD AUC (μm^2^s) with Dyngo4A. Statistical significance determined using a Student’s *t* test.

To quantify changes in the activity-dependent entrapment of Syt1-pH at sites of SV fusion, we directly compared the mobility of Syt1^WT^ -pH and Syt1^K326A/328A^-pH in nerve terminals, taking advantage of the activity-dependent unquenching of Syt1-pH to identify active presynapses. Syt1 mobility was lower compared to that the whole axon (Fig. 1D i-ii) suggesting specific confinement of Syt1 at the presynaptic membrane. We plotted the frequency distribution (%) of the log_10_ diffusion coefficient (D Coeff) [D] (μm^2^ s^-1^) values of Syt1-pH (Fig. 1D i) and the MSD of Syt1-pH (μm^2^) over time (200 ms) (Fig. 1D ii). Although we observed a moderate (p=0.07) shift toward more mobile values for the K326A/K328A mutant (Fig. 1D iii), the MSD of Syt1^K326A/ K328A^-pH was significantly higher than that of Syt1^WT^ -pH (p=0.048) (Fig. 1D iv). This further suggests that the nanoclustering of Syt1-pH at the nerve terminal membrane is controlled via interaction with SV2A. To confirm that the entrapment of Syt1-pH was not a byproduct of internalisation into SVs, uPAINT imaging of Syt1-pH-atto647N-NB was carried out in the presence of Dyngo4A (30 μM for 30 min), a pharmacological agent that blocks the GTPase activity of dynamin, thereby inhibiting endocytosis (McCluskey et al., 2013). Dyngo4A treatment did not alter Syt1 mobility (Fig. 1D v), indicating that the confinement of Syt1 occurring on the PM does not result from dynamin-mediated endocytic events.

### Syt1 nanoclustering at the plasma membrane is impaired by the K326A/K328A mutation

Having established that the K326/K328 residues control the lateral confinement of Syt1-pH in an activity-dependent manner, we next examined the effect of the K326A/K328A mutation on the nanoclustering of Syt1-pH on the PM. For Syt1^WT^ and Syt1^K326A/328A^, nanoclusters were distributed across the axonal branches (Fig. 2A). Dual colour imaging of Syt1-pH-atto647N-NB (uPAINT) was also undertaken in tandem with photoactivated localization microscopy (PALM) of clathrin-mEos4b, revealing that Syt1 nanoclusters were formed in regions absent in clathrin, suggesting that Syt1 nanoclustering at the PM occurs independently of clathrin-mediated SV recycling (Fig. 2B i-iii). Subsequently, we implemented the newly-developed nanoscale spatiotemporal indexing clustering (NASTIC) analysis (Wallis et al., 2021) to define the dimensions of Syt1 nanoclusters (Fig. 2B iv-v). To determine if the increased mobility of Syt1^K326A/K328A^ was due an alteration in nanoclustering, we used NASTIC analysis to quantify the size, density, and apparent lifetime of Syt1^WT^ and Syt1^K326A/K328A^ nanoclusters. We found that a portion of Syt1^WT^-pH molecules (4.68 ± 0.87%) were organized into nanoclusters (0.042 ± 0.0015 μm^2^) of short duration (6.18 ± 0.28 sec). In contrast, Syt1^K326/328A^-pH nanoclustering was less prominent (Fig. 2C i-ii). The MSD (Fig. 2D i), detection frequency (Fig. 2D ii), and apparent lifetime (Fig. 2D iii) of clustered Syt1 was unaffected by K326/328A. However, the K326A/K328A mutant caused a significant loss in nanocluster density (p=0.048) (Fig. 2D iv), and a corresponding increase in both cluster area (p=0.012) (Fig. 2D v) and radius (p=0.008) (Fig. 2D vi). Our results demonstrate that binding to SV2A (through the K326/K328 residues) is critical for Syt1 nanocluster formation during stimulation.

**Figure 2.**
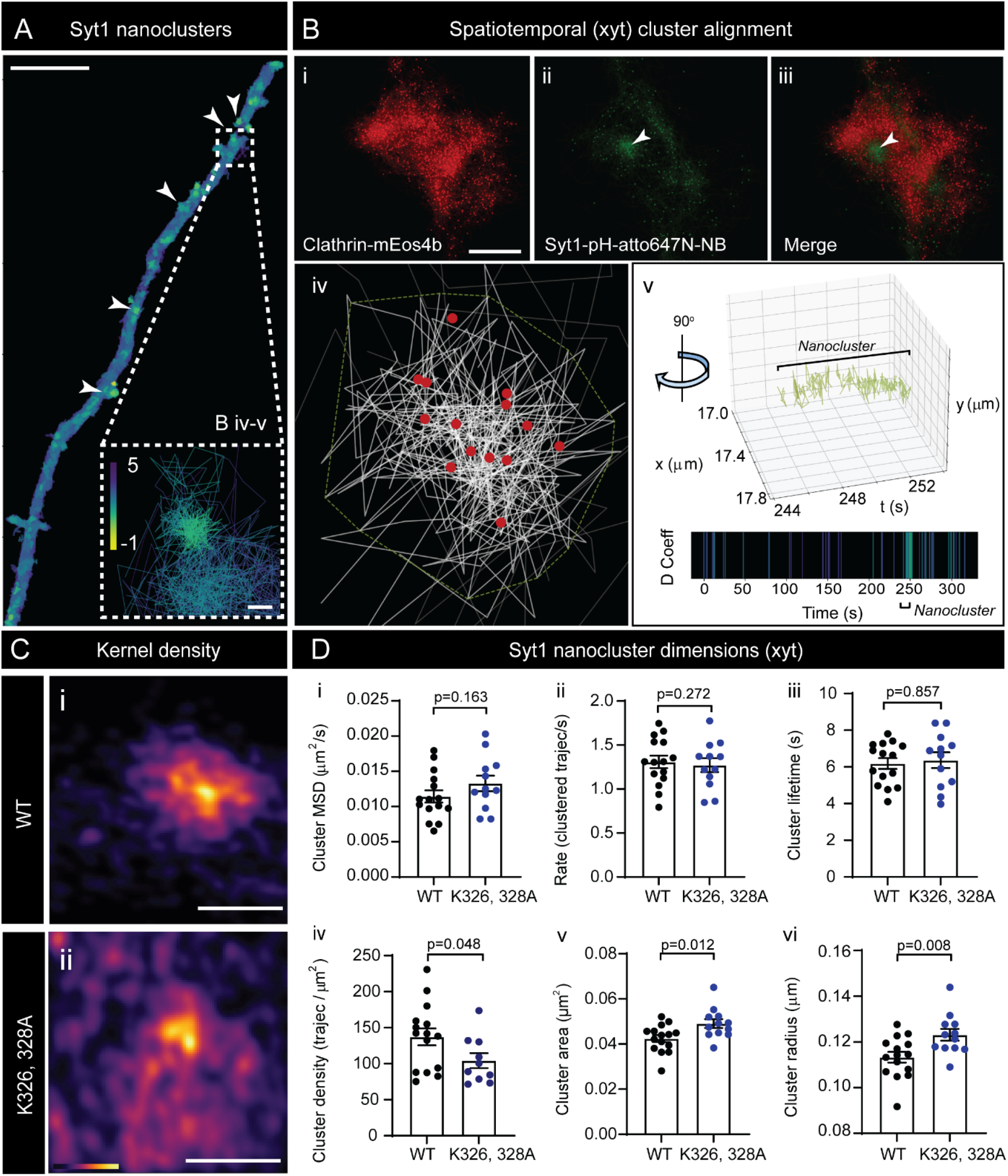
The Syt1 K326A/K328A mutation decreases the density and increases the size of Syt1 nanoclusters. **(A)** Syt1-pH-atto647N-NB PM nanoclusters, indicated by white arrows (scale bar = 5 μm). Insert shows example nanocluster trajectories (analysed in B iv-v) with variable D Coeff values, (scale bar = 0.1 μm). **(B)** Spatiotemporal alignment (xyt) of Syt1^WT^ nanoclusters in presynapse. (i) Clathrin-mEos4b and (ii) Syt1-pH-atto647N-NB trajectories were (iii) merged to show segregation of Syt1 nanoclusters (green) from clathrin-mEos4b (red). (iv) Enhanced view of Syt1 tracks shown in (A) converging within a single nanocluster with nanocluster boundary (green outline) and centroids (red) shown. (v) Temporal longevity of clusters shown in B iv (tracks rotated ~90° across xyt) obtained using Nanoscale Spatiotemporal Indexing Clustering (NASTIC). **(C)** Kernel Density Estimate (KDE) (i) Syt1^WT^-pH and (ii) Syt1^K326A/K328A^-pH. Scale bar = 0.2 μm. **(D)** Syt1^WT^ and Syt1^K326A/K328A^ cluster dimensions. Nanoclustered trajectory (i) MSD (μm^2^ for 200 ms) (ii) Rate (trajec/s) (iii) lifetime (iv) density (trajectories/μm^2^) (v) area (μm^2^) (vi) radius (μm). Statistical significance determined using a Student’s *t*-test.

### Knockdown of endogenous SV2A increases the surface mobility of Syt1 and impairs Syt1 nanocluster formation

To confirm that the increased surface mobility of Syt1 caused by the K326A/K328A mutation was specific to altered SV2A binding, we transfected neurons with an SV2A-shRNA tagged with mCerulean (mCer) to knockdown the endogenous population of SV2A (Fig. 3A) (Dong et al., 2006; Harper et al., 2020; Zhang et al., 1994; Zhang et al., 2015). As previously reported, a significant decrease in endogenous SV2A expression was observed in the presence of SV2A-shRNA-mCer (Fig. 3A). Therefore, we performed uPAINT imaging and tracked the mobility of Syt1-pH-atto647-NB in nerve terminals in the presence of either mCer, SV2A-shRNA-mCer. To further validate the effect of SV2A on Syt1-pH mobility, we also performed a rescue experiment in which we co-expressed SV2A-shRNA with shRNA-resistant SV2A-mCer and Syt1-pH in neurons (Fig. 3B-C). As expected, knockdown of endogenous SV2A with SV2A-shRNA-mCer increased the surface mobility of Syt1-pH at the presynapse, leading to a significant decrease in the percentage of immobile Syt1-pH molecules (Fig. 3C i) and of the MSD AUC (Fig. 3C ii-iii). Importantly, the mobility of Syt1-pH-atto647N-NB was rescued upon co-expression SV2A-mCer compared to SV2A shRNA-knockdown alone (Fig. 3C i-iii).

**Figure 3.**
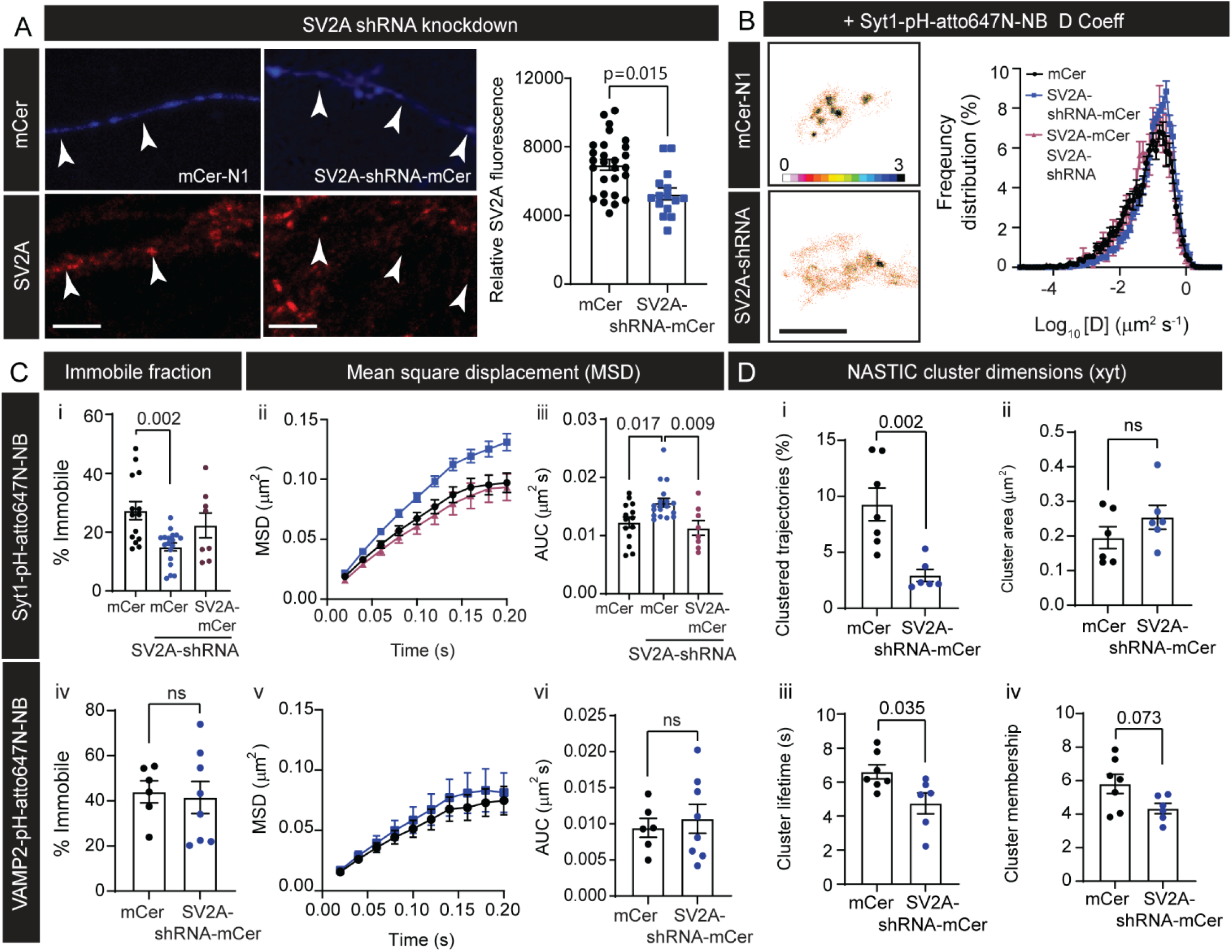
SV2A depletion increases Syt1 mobility which is rescued by SV2A-mCer re-expression. **(A)** Expression of mCer or SV2A-shRNA-mCer with endogenous SV2 staining. Arrows highlight transfected axons. A significant decrease in SV2 was observed in the presence of SV2A-shRNA (p=0.015) **(B)** Intensity map of Syt1-pH-atto647N-NB+mCer and Syt1-pH-atto647N-NB+SV2A-shRNA-mCer are shown together with frequency distribution of Log_10_ [D] (μm^2^s^-1^) values for Syt1-pH-atto647N-NB trajectories. **(C)** (i-iii) Syt1 mobility in the presence of mCer, SV2A-shRNA-mCer or SV2A-mCer-SV2A-shRNA: (i) % immobile (ii) MSD (μm^2^ over 200 ms) and (iii) MSD AUC (μm^2^s). VAMP2 mobility in the presence of mCer or SV2A-shRNA-mCer: (iv) % immobile, (v) MSD (μm^2^ over 200 ms), and (vi) AUC (μm^2^s). **(D)** Syt1 nanoclustering following SV2A knockdown (i) clustered trajectories (%). (ii) Area (μm^2^) (iii) membership (trajec/cluster) (iv) lifetime (s) of nanoclusters. A significant decrease in % clustered trajectories (p=0.002), cluster area (p=0.008) and lifetime (p=0.035) was observed in the presence of SV2A-shRNA knockdown. Statistical significance determined using a one-way ANOVA and Tukey’s test for multiple comparisons and Student’s *t*-test for single comparisons.

Based on the increased mobility of Syt1-pH-atto647N-NB following SV2A knockdown, we concluded that SV2A plays a key role in sequestering Syt1 into nanoclusters at the PM. However, it is not clear whether this function is specific to Syt1. To determine whether SV2A helps sequester other vesicular machinery on the PM, we performed uPAINT imaging of VAMP2-pHluorin (VAMP2-pH) in the presence of SV2A-shRNA-mCer. The mobility of VAMP2-pH-atto647NB remained unaffected following SV2A knockdown, with no observed shift in the percentage of immobile trajectories (Fig. 3C iv) and MSD (Fig. 3C v-vi). Although we cannot rule out that other proteins play a role in trapping Syt1 on the PM, our results suggest that the entrapment effect of SV2A is limited to Syt1. Finally, we examined the impact of SV2A knockdown on Syt1 nanoclustering using NASTIC. Expression of SV2A-shRNA caused a reduction in the percentage of clustered trajectories at the PM (p=0.002) (Fig. 3D i) and decrease in Syt1 nanocluster area (p=0.006) (Fig. 3D ii), although no reduction in the molecular occupancy of these clusters was observed (Fig. 3C iii). Further, the apparent lifetime of Syt1 nanoclusters was also significantly decreased in the presence of SV2A-shRNA-mCer (p=0.035) (Fig. 3D iv), likely accounting for the increased surface mobility of Syt1 when SV2A is downregulated.

### SV2A mobility is dependent on Syt1 binding, but not on interaction with the clathrin adaptor AP2 or dynamin

Our results demonstrate that the nanocluster organisation of Syt1 is dependent on its interaction with SV2A. We therefore investigated whether SV2A nanoscale organisation was reversibly controlled by Syt1. To this end, we used a mutant of the cognate interaction site (SV2A-T84A) and examined its nanoscale mobility (Fig. 4A). Additionally, to determine whether clathrin endocytic machinery could affect SV2A nanoclustering, we examined the effect of inhibiting SV2A interaction with the clathrin adaptor AP2 using an established mutant (SV2A^Y46A^) (Yao et al., 2010) (Fig. 4A). To confirm the effect of SV2A^T84A^ on Syt1 binding, we coimmunoprecipitated SV2A from HEK293 cells expressing both HA-Syt1^WT^ and either SV2A^WT^-mCer or SV2A^T84A^-mCer (Fig. 4B). The binding was significantly reduced by the T84A mutation (Fig. 4B i, ii) (Zhang et al., 2015). Interestingly, the Y46A mutant significantly increased the amount of Syt1-HA pulldown (Fig. 4B i-ii). This finding suggests that AP2 and Syt1 compete for interaction with SV2A, likely due to the proximity of the Syt1 (T84) and AP2 (Y46) interaction sites (Fig. 4A). Next, uPAINT imaging of hippocampal neurons expressing SV2A-pHluorin (SV2A-pH) was performed (Fig. 4C-D). SV2A-pH formed nanoclusters within presynaptic boutons, which are less prominent in the presence of the T84A mutation (Fig. 4C). This was due to a decrease in the apparent lifetime of SV2A^T84A^ nanoclusters (Fig. 4C i). Nanocluster area and membership were unchanged (Fig. 4C ii, iii). SV2A-pH mobility was significantly increased in the presence of the T84A mutation (Fig. 4D i-ii) demonstrating that SV2A-Syt1 interaction is essential for the trapping of both molecules at the PM. However, the loss of AP2 binding had no effect on the surface mobility of SV2A (Fig. 4D i-ii), suggesting that nanoclustering of SV2A is not regulated by the endocytic machinery. No change in the MSD of SV2A was observed following Dyngo4A (30 μM for 30 min)-induced dynamin inhibition (Fig. 4D iii-iv), further confirming this observation. Overall, our results suggest that the nanoclustering of Syt1 and SV2A is controlled by their intramolecular interaction.

**Figure 4.**
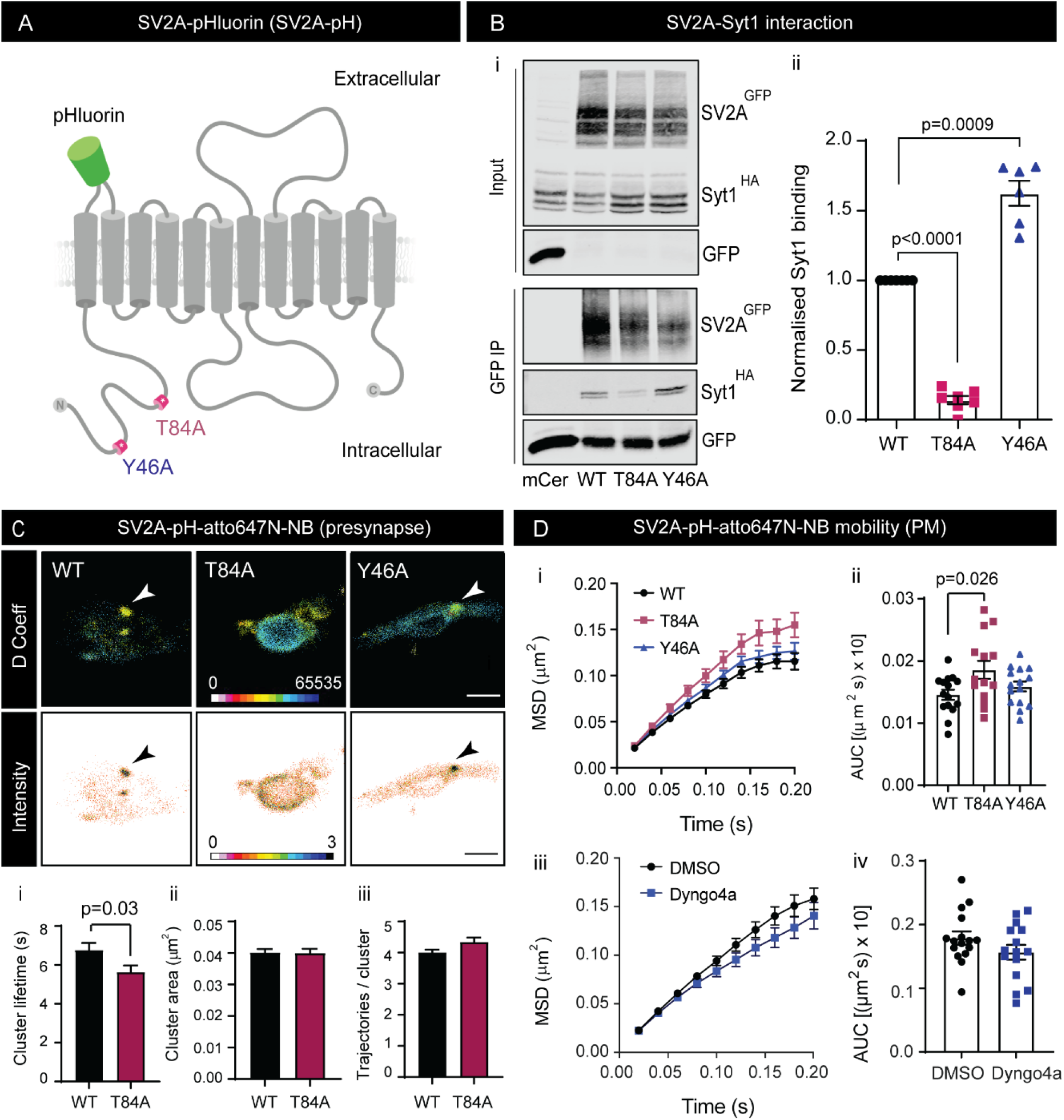
SV2A nanoclustering is controlled by Syt1 interaction (T84A) and unaffected by AP2 interaction (Y46A) and dynamin inhibition. **(A)** Structure and orientation of SV2A-pH across PM. pH tag (green) is located between first and second transmembrane domains. Note the proximity of the T84A and Y46A epitopes (red). **(B)** Co-IP of SV2A-mCer with Syt1-HA. (i) Representative blots from total protein input and GFP IP (ii) Normalised Syt1-HA binding to SV2A-mCer (WT, T84A and Y46A). **(C)** SV2A (WT, T84A or Y46A)-pH nanoscale organization within the presynapse of hippocampal nerve terminals. For D Coeff panels, regions highlighted in warm colours represent points of low mobility. Arrows indicate nanoclusters points. SV2A nanoclustering in the presence and absence of Syt1 interaction (T84A) as determined by NASTIC. (i) Nanocluster lifetime (s) (ii) area (μm^2^) and (iii) membership (trajectories per cluster). **(D)** Surface mobility of SV2A^WT^, SV2A^T84A^ and SV2A^Y46A^ at the presynapse. (i) MSD (μm^2^) of SV2A-pH-atto647-NB over time (200 ms) (ii) MSD AUC (μm^2^ s × 10) (iii) SV2A^WT^-pH-atto647N-NB MSD following Dyngo4A treatment (30 μM for 30 min) (iv) AUC (μm^2^ s x 10). Scale bar = 1 μm. Statistical significance determined by a one-way ANOVA with a Tukey’s test for multiple comparisons and a Students’ *t*-test for single comparisons.

### Activity-dependent retrieval of SV2A is controlled by AP2 but not Syt1

To determine whether the clustering of SV2A is associated with alterations in SV endocytosis, we examined the kinetics of SV2A-pH retrieval. SV2A-pH (WT, T84A, Y46A) was expressed in hippocampal neurons and the fluorescence decay of pH was examined following electrical field stimulation (300 action potentials (APs) at 10 Hz), prior to treating neurons with ammonium chloride (NH_4_Cl) to reveal the total population of SV2A-pH (Fig. 5A). The loss of pH fluorescence is reflective of the kinetics of its retrieval during SV endocytosis, since this is rate limiting in comparison to subsequent SV acidification (Atluri and Ryan, 2006; Granseth et al., 2006). The retrieval of the Y46A mutant was compromised, suggesting that interactions with AP2 are required for its efficient recovery during endocytosis (Fig. 5B i-ii). However, impairing the interaction of SV2A-pH with Syt1 (SV2A^T84A^) did not impact SV2A retrieval (Fig. 5B i-ii) (Zhang et al., 2015). The SV2A^Y46A^ mutant retrieval delay was not due to alterations in SV exocytosis, since no difference in the evoked fluorescence peak was observed as a proportion of the total population of SV2A-pH, which was also the case for the SV2A^T84A^ mutant (Fig. 5B iii). These findings demonstrate that AP2-mediated endocytosis of SV2A-pH occurs independently of SV2A-Syt1 interaction and nanoclustering.

**Figure 5.**
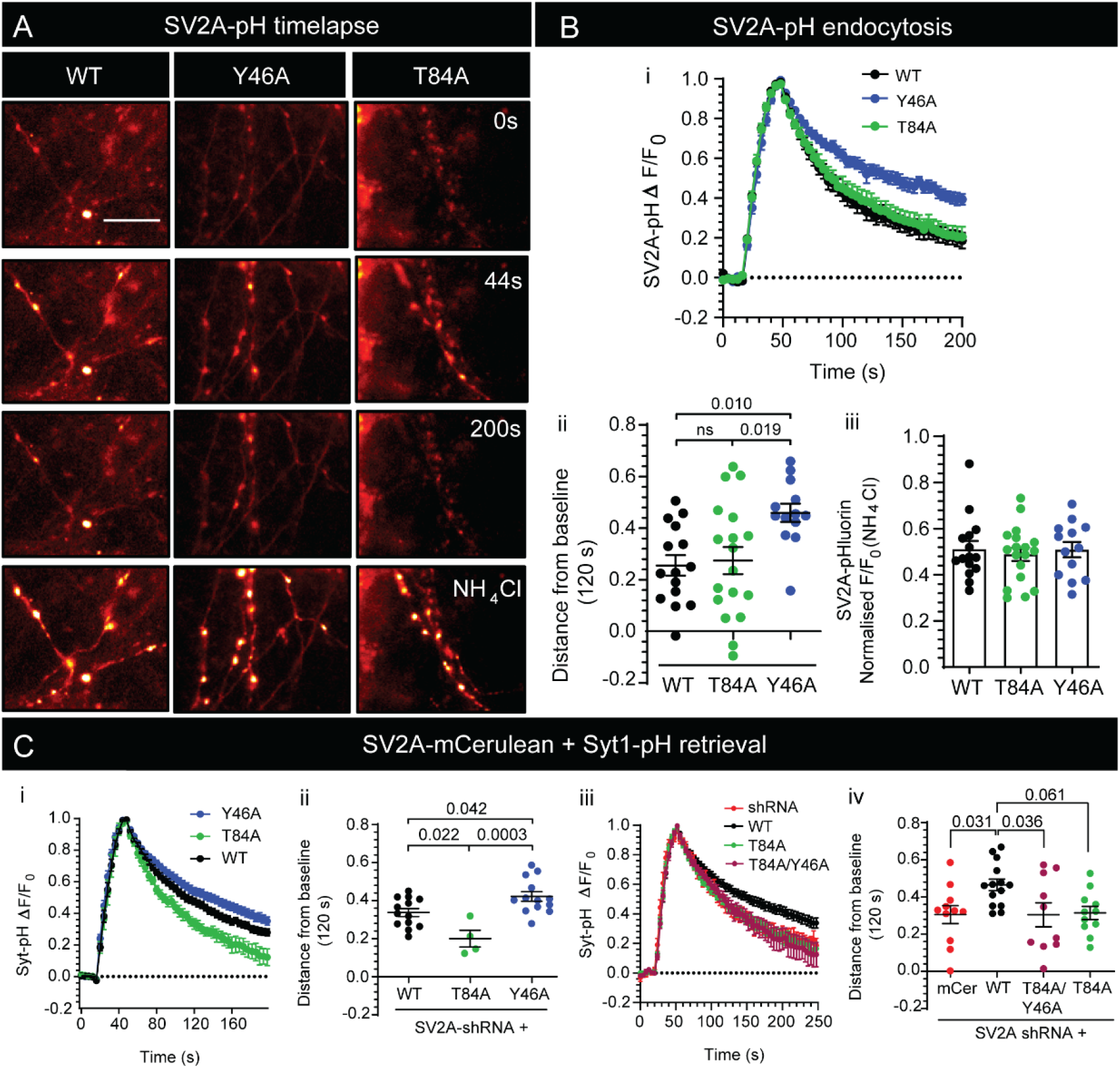
Activity-dependent endocytosis of Syt1-pH is controlled by SV2A. **(A)** SV2A^WT^, SV2A^T84A^ and SV2A^Y46A^ -pH surface fluorescence time-lapse following stimulation (10Hz, 300 AP), prior to treatment with ammonium chloride (NH_4_Cl). Scale bar = 20 μm. **(B)** Loss of AP2 binding (Y46A) impairs SV2A-pH retrieval. (i) SV2A-pH retrieval time course (WT, T84A, Y46A). (ii) SV2A-pH remaining to be retrieved (distance from baseline (at 120 s)). (iii) Normalised F/F_0_ (NH_4_Cl perfusion). **(C)** Syt1-pH activity-dependent retrieval kinetics in SV2A knockdown neurons in the presence SV2A-mCer expression. (i) Time course of Syt1-pH fluorescence in the presence of SV2A^WT^, SV2A^T84A^ and SV2A^Y46A^-mCer after stimulation (10Hz, 300AP) quantified by (ii) distance from baseline (120 s). (iii) Time course of Syt1-pH fluorescence in the presence of SV2A^WT^, SV2A^T84A^ and SV2A^T84A/Y46A^ following stimulation (10Hz, 300AP) quantified with (iv) distance from baseline (120s). Statistical significance determined with Kruskal-Wallis test with Dunn’s test for multiple comparisons.

### SV2A controls the activity-dependent endocytosis of Syt1

Knockdown of SV2A and disruption of binding of Syt1 to SV2A accelerates the internalisation of Syt1-pH during SV endocytosis, suggesting it may retard this process (Harper et al., 2020; Kaempf et al., 2015; Zhang et al., 2015). To determine whether the retrieval of Syt1-pH was dependent on SV2A, we performed molecular replacement experiments in SV2A-depleted neurons with wild-type and mutant variants of SV2A. Hippocampal neurons were cotransfected with a bicistronic plasmid expressing SV2A-shRNA and Syt1-pH with coexpression of either SV2A^WT^-mCer, SV2A^T84A^-mCer (Syt1 binding mutant), or SV2A^Y46A^-mCer (AP2 binding mutant). We confirmed that disruption of SV2A-Syt1 interaction via SV2A^T84A^-mCer accelerated the retrieval kinetics of Syt1-pH from the PM, compared to SV2A^WT^-mCer (Fig. 5C i-ii) (Zhang et al., 2015). Conversely, blocking the interaction between SV2A and AP2 with the Y46A mutant of SV2A, slowed the activity-dependent retrieval of Syt1-pH (Fig. 5C i-ii). However, the accelerated retrieval of Syt1-pH caused by the expression of SV2A^T84A^ was not rescued by the additive loss of AP2 binding (SV2A^T84A/Y46A^) (Fig. 5C iii-iv), indicating that SV2A interaction is the dominant mechanism by which Syt1 endocytosis is regulated.

### Loss of SV2A interaction alters the intracellular sorting of Syt1 at the recycling pool of SVs

The accelerated retrieval of Syt1 caused by SV2A^T84A^ raises the question of whether SV2A controls the endocytic targeting of Syt1 to recycling SVs. This suggests that interfering with SV2A causes intracellular mis-sorting of Syt1. To determine whether SV2A controls the nanoscale organization of internalised Syt1-pH, a sub-diffractional Tracking of Internalised Molecules (sdTIM) technique SVs (Joensuu et al., 2017; Joensuu et al., 2016) was used to image endocytosed Syt1. Syt1-pH-positive neurons were pulsed (56 mM K^+^ with anti-GFP-atto647N-NB for 5 min) before being washed and chased under resting conditions (5.6 mM K^+^for 5 min). To selectively image the recycling pool of SVs containing Syt1 with greater accuracy, we digested Syt1-pH-atto647N-NB at the Tobacco-Etch Virus (TEV) cleavage sequence present between Syt1 and the pH tag, using an active TEV (AcTEV) protease (1 mM, 15 min) (Fig. 6A i-iii) (Gimber et al., 2015; Hua et al., 2011; Nair et al., 2013; Wienisch and Klingauf, 2006). Accordingly, we observed a significant decrease in the fluorescence of Syt1^WT^-pH and Syt1^K326A/K328A^-pH in the presence of active (AcTEV) protease, compared to the inactivated, boiled AcTEV control (95°C for 10 min) (Fig. 6A iv-v).

**Figure 6.**
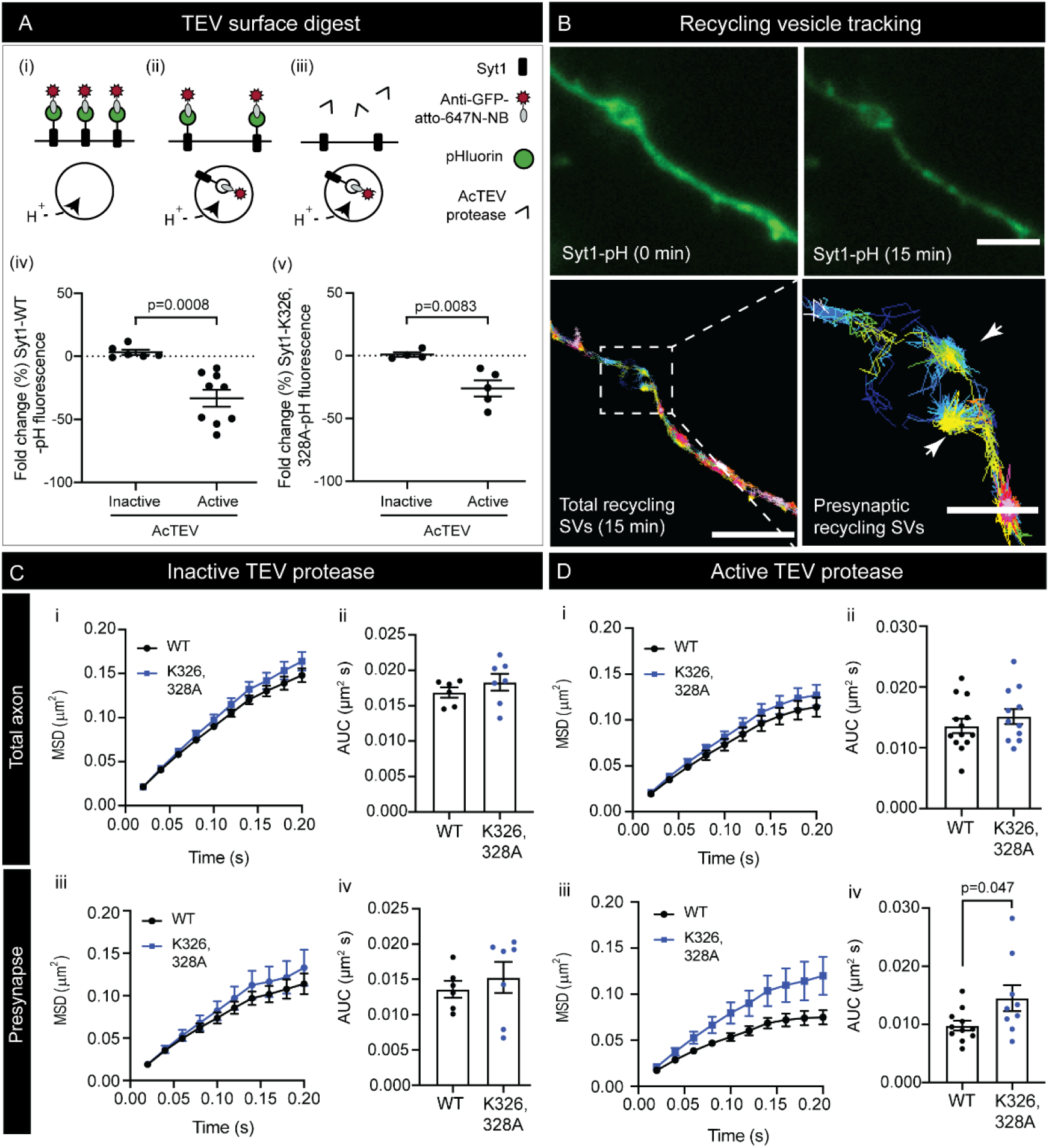
Sub-diffractional tracking of internalised molecules (sdTIM) of Syt1-pH reveals an alteration in the intracellular sorting of Syt1 in the absence of SV2A interaction. **(A)** Digestion of the surface fraction of Syt1-pH-atto647N-NB with AcTEV protease. Method overview: (i) Atto647N-NB binds Syt1-pH during stimulation (56mM K^+^ for 5 min), (ii) Syt1-pH-atto647N-NB is internalised following a chase step (5.6 mM K^+^; 5 min) and (iii) TEV digest, which removes pH-atto647N-NB. (iv-v) Fold change (%) in Syt1-pH fluorescence in the presence of the inactive (boiled) and active AcTEV protease for (iv) Syt1^WT^-pH and (v) for Syt1^K326A/K328A^-pH. **(B)** Loss of Syt1-pH fluorescence following a 15-minute incubation with AcTEV protease to cleave the surface fraction of Syt1-pH-atto647N-NB and track Syt1-pH-atto647N-NB within the recycling pool of SVs. Scale bar = 4 μm (axon) and 2 μm (presynapse). **(C-D)** Mobility of Syt1^WT^- and Syt1^K326A/K328A^-pH-atto647N-NB in the presence of **(C)** inactive and **(D)** active TEV: (i-ii) Total axon and (iii-iv) Presynaptic MSD (μm^2^ over 200 ms) and AUC (μm^2^s). A significant increase in the displacement of Syt1^K326A/K328A^-pH-atto647N-NB (p=0.047, Student’s t-test) was detected at the presynapse upon digest of the surface fraction of Syt1-pH-atto647N-NB using AcTEV.

Following internalisation of the surface population, recycling SVs containing Syt1-pH-atto647N-NB were tracked across the total axon and in nerve terminals (Fig. 6B). A control experiment was carried out using boiled, inactivated protease (Fig. 6C). In the absence of surface digestion, Syt1^WT^-pH and Syt1^K326A/K328A^-pH had a comparable mobility across the entire axon (Fig. 6C i-ii) and within nerve terminals (Fig. 6C iii-iv). However, upon digestion of the surface fraction of Syt1-pH with the active protease, Syt1 mobility was different (Fig. 6D). Specifically, the MSD of Syt1-pH-atto647N-NB was lower in the presence of the active protease compared to the inactive protease, indicating entry of Syt1 into the recycling pool of SVs (Fig. 6C-D) (Joensuu et al., 2016). Syt1^WT^-pH and Syt1^K326A/K328A^-pH had similar mobilities across the whole axon (Fig. 6D i-ii). Surprisingly, Syt1^K326A/K328A^-pH was significantly more mobile at the presynapse (Fig. 6D iii-iv). These results suggest that loss of SV2A binding causes intracellular mis-sorting of Syt1 in nerve terminals. It is unlikely that these differences are due to stranding of Syt1 at the PM, as no significant differences were observed without surface fraction digestion. Therefore, loss of SV2A binding causes entry of Syt1 to a different endocytic compartment leading to differences in intracellular sorting.

### Loss of SV2A interaction alters Syt1 trafficking away from Rab5-endosomes

The elevated mobility of Syt1^K326A/K328A^-pH following endocytosis within nerve terminals suggests that reduced binding to SV2A causes mis-sorting of Syt1 into a more mobile endocytic compartment. Syt1 has previously been shown to be localized in early and recycling endosomes following internalisation 20-minutes post-fusion (Diril et al., 2006). It is becoming increasingly apparent that at physiological temperatures, clathrin-dependent and –independent cargo sorting occurs at the level of internalised endosomes (Ivanova et al., 2021; Kononenko et al., 2013; Watanabe et al., 2014). Therefore, the differences in Syt1-pH mobility observed upon internalisation may stem from sorting back to the recycling SV pool or the endolysosomal system. To address this, we examined the co-localisation of internalised Syt1 with Rab5, a GTPase associated with early endosomes (Bucci et al., 1992) and bulk endosomes (Kokotos et al., 2018). Synaptotagmin has previously been shown to cluster within Rab5-positive early endosomes following internalisation (Hoopmann et al., 2010). Neurons were co-transfected with either Syt1^WT^-pH or Syt1^K326A/K328A^-pH and Rab5-mRFP (Vonderheit and Helenius, 2005). Following activity-dependent internalisation of Syt1-pH-atto647N-NB, 3D structured illumination microscopy (3D-SIM) was carried out to determine the level of Syt1 trafficking to Rab5-positive early endosomes (Fig. 7A-B). We defined populations of Rab5-mRFP clusters as surfaces that encased internalised molecules of Syt1-pH-atto647N-NB (Fig. 7C). A higher proportion of Syt1^WT^-pH-atto647N-NB localizations was identified in 3D-generated Rab5 surfaces (p=0.03) compared to Syt1^K326A/K328A^-pH-atto647N-NB (Fig. 7D). The volumetric density of Syt1^WT^-pH-atto647-NB within Rab5-mRFP surfaces was also significantly higher (p=0.02) compared to the K326A/K328A mutant (Fig. 7E). No significant difference in the endosomal surface volume was observed between wildtype and mutant Syt1 (Fig. 7F). Based on these findings, we concluded that inhibiting SV2A interaction altered Syt1 intracellular trafficking towards Rab5-positive compartments, elevating Syt1’s intracellular mobility (Fig. 8A-E).

**Figure 7.**
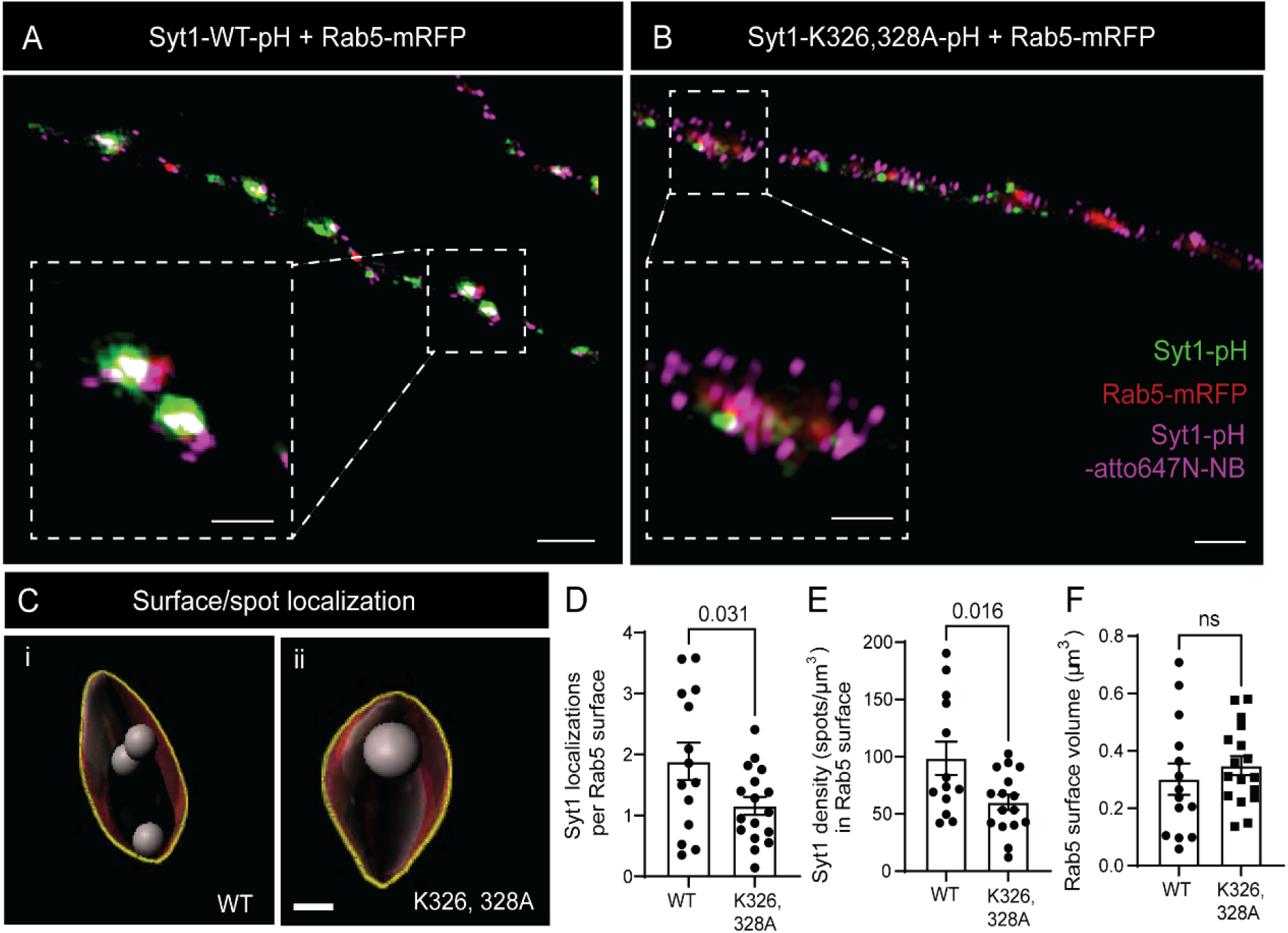
SV2A interaction elevates trafficking of Syt1 to a Rab5-positive endocytic pathway. 3D-structural illumination microscopy of neurons transfected with Rab5-mRFP and **(A)** Syt1^WT^-pH **(B)** Syt1^K326A/K328A^ -pH. **(C)** Surfaces generated from clusters of Rab5-mRFP immunofluorescence and corresponding spots of Syt1^WT^ -pH and Syt1^K326A/K328A^ -pH- atto647N-NB within each surface. Spots (atto647N-NB) are depicted in grey. Outline of surfaces (Rab5-mRFP clusters) are shown in yellow. **(D)** Syt1-pH-atto647N-NB localizations per Rab5 surface. **(E)** Syt1-pH-atto647N-NB density (#spots/μm^3^) within Rab5-mRFP surfaces. **(F)** Rab5-mRFP surface volume (μm^3^). Statistical differences determined using Student’s t test. Scale bar = 2 μm.

**Figure 8.**
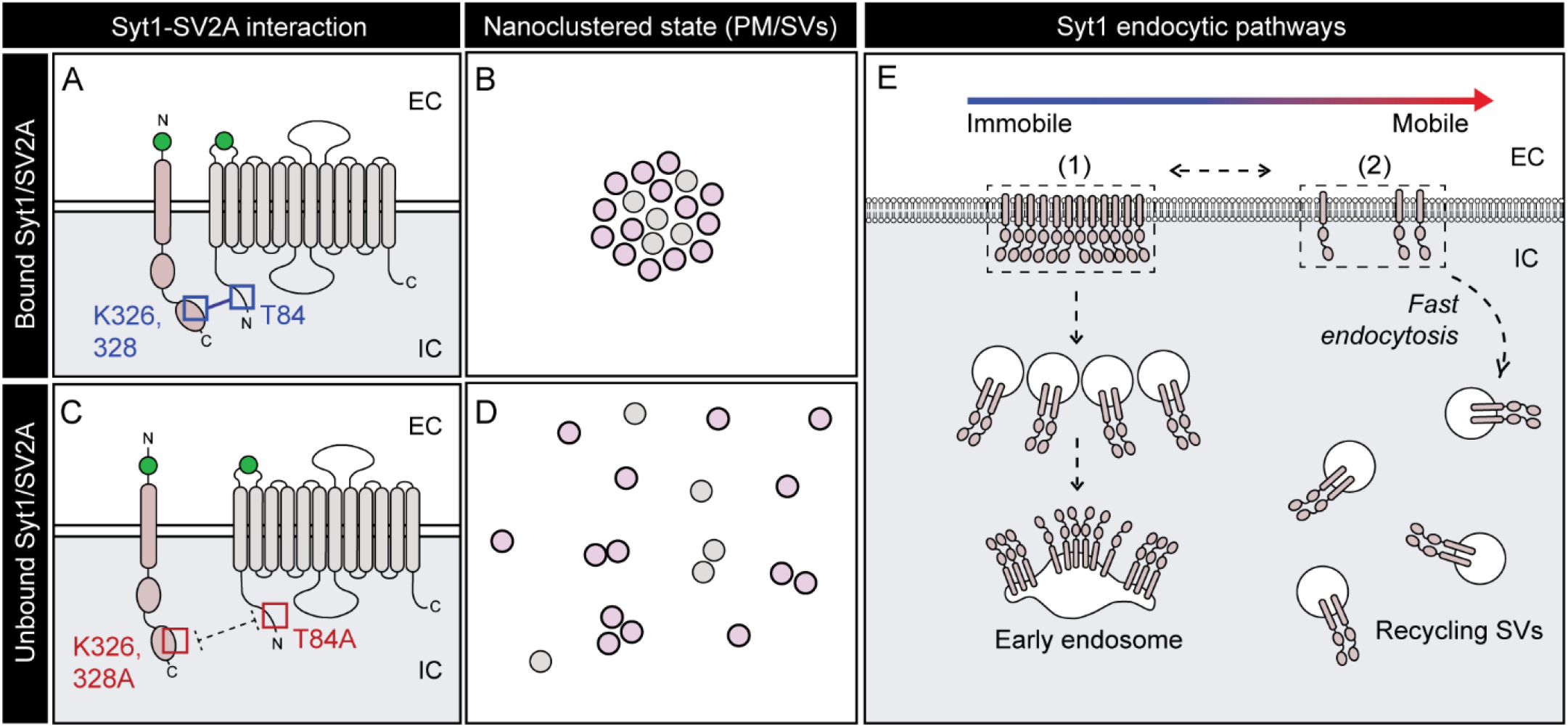
Hypothetical model: SV2A-bound Syt1 nanoclusters control pathway of Syt1 endocytosis causing changes in intracellular sorting. **(A)** The SV2A-Syt1 complex linked by the K326/K328 epitopes at the polybasic region of the C2B domain and the T84 epitope at the N-terminal tail of SV2A. **(B)** Schematic of Syt1 and SV2A grouped together in nanoclusters at the PM. **(C)** Unbound Syt1 and SV2A due to loss of binding (K326A/K328A or T84A). **(D)** Schematic of unbound Syt1 and SV2A with increased surface displacement. **(E)** Syt1 recycling pathways (1) Early endosome recruitment (2) Recruitment into recycling SVs. Colour gradient represents changes in surface mobility.

## Discussion

Interactions between vesicular proteins on the PM help maintain the organization, stoichiometry, and composition of SVs by enhancing the fidelity of endocytic events (Gordon and Cousin, 2013). In this study, we provide evidence that their nanoclustering at the PM controls the targeting of vesicular machinery into recycling SVs. We demonstrate that SV2A controls the nanoclustering of Syt1 at the PM through interactions of its cytoplasmic N-terminus with the K326/328 epitopes within the polybasic region of the C2B domain of Syt1. Importantly, this mechanism also works in reverse: SV2A is also sequestered into nanoclusters through its interaction with Syt1. This dual entrapment underpins the formation of nanoclusters at the PM. The endocytic machinery (AP2, dynamin) does not play a role in this process during the timeframe of our experiments. Furthermore, the interaction between Syt1 and SV2A delays the kinetics of Syt1 retrieval (Harper et al., 2020; Kaempf et al., 2015; Zhang et al., 2015) and may control entry of Syt1 into discrete intracellular compartments.

### Mechanisms underpinning SV2A-Syt1 nanoclustering

The finding that SV2A controls Syt1 surface nanoclustering through interaction with the K326/K328 residues is in accordance with previous observations which showed that substituting the K326, 328 epitopes with an alanine decreases Syt1 oligomerization (Chapman et al., 1998). SV2A nanoclustering is also regulated by cognate interaction with Syt1. This is similar to other presynaptic molecules, such as syntaxin1A (Bademosi et al., 2016; Lang et al., 2001; Sieber et al., 2006) and Munc18 (Kasula et al., 2016), which form nanoclusters dependent on molecular interactions and play a key role in exocytosis. Our results demonstrate that SV2A-Syt1 nanoclustering is not directly regulated by AP2 and dynamin and is likely independent of the endocytic machinery. Importantly, VAMP2 forms surface nanoclusters that are controlled by the endocytic machinery (Gimber et al., 2015), raising the possibility that several layers of clustering mechanisms take place at the PM. Vesicular proteins are therefore pre-assembled on the PM with various levels of control, either by direct binary interaction or via the endocytic machinery, thereby controlling the fidelity of uptake into recycling SVs. This finding is supported by the observation that Syt1 and AP2 compete for interaction with SV2A, and that the endocytosis kinetics of SV2A are unaffected by Syt1 interaction (T84A), but dependent on AP2 binding (Y46A). Syt1 may be more accessible to the endocytic machinery in the absence of SV2A binding, as the absence of SV2A promotes the uptake of Syt1-pH into recycling SVs, while nanocluster formation is isolated from clathrin and unaffected by dynamin inhibition. Furthermore, the lack of bidirectional control of the retrieval of Syt1 and SV2A may be due to both molecules engaging different sets of endocytic adaptor molecules, which may lead to differing patterns of surface nanoclustering. For example, Syt1 has a unique interaction with stonin 2, an endocytic adaptor that appears to exclusively shepherd Syt1 into clathrin-coated pits (CCPs) in concert with AP2 (Diril et al., 2006). Loss of stonin 2 accelerates endocytosis of Syt1 (Kononenko et al., 2013), a phenotype mimicked by SV2A knockdown. Examining the effect of stonin 2 on Syt1 nanoclustering will be an important area of interest for future studies.

### SV2A-mediated nanoclustering promotes Syt1 trafficking to Rab5-positive early endosomes

Our results demonstrate that the nanoclustering of Syt1 by SV2A at the PM controls Syt1 internalisation. This occurs either by delaying entry into recycling vesicles, or by dictating the intracellular targeting into recycling SVs. Release of SV2A from nanoclusters is required for AP2-mediated sorting, as SV2A nanoclustering hinders interaction with the endocytic machinery. This may involve steric limits, as the nanoclusters quantified in our analysis are significantly larger than CCPs (0.065-0.125 μm) (Kirchhausen and Harrison, 1981) and recycling SVs (0.040 μm in diameter) (Zhang et al., 1998). Importantly, the Syt1-SV2A interaction, and by extension nanoclustering, depends on casein kinase phosphorylation of the T84 epitope (Zhang et al., 2015). Thus, the SV2A-Syt1 nanoclustering is likely a highly regulated process. Surface nanoclustering may act as a transition state for Syt1 at the PM and allow neurons to fine-tune endocytosis, titrating the internalisation of Syt1 back into recycling SVs. As such, it will be important to determine the conditions during which phosphorylation of SV2A’s N-terminal domain (T84) occurs, and if this process occurs at the PM. Importantly, Syt1 has been shown to control endocytic events (Li et al., 2017; Nicholson-Tomishima and Ryan, 2004; Poskanzer et al., 2003) and recruit endocytic machinery during endocytosis in endocrine cells (McAdam et al., 2015). The C2 domains of Syt1 regulate the kinetics of vesicle internalisation in a calcium-dependent manner, similar the action of the C2 domains during exocytosis (Yao et al., 2012). SV2A binding to the C2B domain of Syt1 is negatively regulated by interaction of the C2B domain with Ca^2+^ (Schivell et al., 2005). Therefore, it is tempting to speculate that the pace of Syt1-mediated endocytosis may be decreased through competitive interaction with Ca^2+^ and nanoclustering by SV2A.

While surface nanoclustering restricts the pace of Syt1 endocytosis, it is also likely to regulate internalisation via alternative modes of recruitment. Surface SV2A-Syt1 nanoclusters may form a reservoir controlling entry into recycling SVs, in which smaller subdomains of SV2A-Syt1 are internalised in unison at a slow pace into select endocytic compartments. We demonstrate that SV2A-Syt1 interaction controls the intracellular sorting of Syt1 to Rab5-positive endosomes. Therefore, Syt1 nanoclustering likely restricts access to smaller endocytic pits and facilitates endocytosis into a more mobile pool of recycling SVs, while simultaneously acting as a reservoir for recruitment into Rab5-positive recycling SVs preferentially sorted into early endosomes (Wucherpfennig et al., 2003). This may occur to rescue stranded SV2A-Syt1 during repetitive rounds of SV fusion to avoid nanocluster build up at the PM. Targeting to Rab5-positive endosomes was shown to facilitate intermixing of molecular cargoes with the readily releasable pool (RRP) of SVs, with inhibition of Rab5-mediated endosomal sorting causing reductions in RRP size (Hoopmann et al., 2010). Another possibility is that clustered Syt1-SV2A stranded at the PM is internalised via activity-dependent bulk endocytosis (ADBE). Bulk endosomes, which are positive for Rab5 (Kokotos et al., 2018) and large enough to engulf Syt1 nanoclusters, have been shown to act as a sorting station for SV cargoes in which proteins can be re-routed back to the reserve pool of SVs (Cheung et al., 2010) or trafficked to the endolysosomal system (Ivanova et al., 2021). Therefore, ADBE of SV2A-Syt1 during sustained neurotransmission may act as an intermediate step to the reformation of SVs.

### SV2A-Syt1 surface nanocluster impact on neurotransmission

The role of the SV2 family of proteins in neurotransmission is not well understood. Studies suggest that SV2A controls the size of the RRP (Custer et al., 2006), primes Ca^2+^-dependent release and short-term synaptic plasticity (Chang and Südhof, 2009). The nanoclustering of SV2 and Syt1 may therefore play a key role in neurotransmitter release and plasticity. Understanding the molecular steps involved in this process provides insight into the role of SV2A and Syt1 during synaptic dysfunction, neurological disorders and avenues for therapeutic treatment. Our findings demonstrate that SV2A interaction with Syt1 mediates their nanoclustering at the PM which in turns control their fate in the recycling SVs process. Furthermore, SV2 acts as a gateway for the entry of botulinum neurotoxin type-A (BoNT/A) into neurons. This process is highlighted in a related study, which demonstrates that BoNT/A hijacks these nanoclusters to promotes its internalisation in SVs highlighting their critical importance in vesicular targeting (Joensuu et al., accompanying manuscript).

It has been suggested that interactions between vesicular proteins on the PM help maintain the organization, stoichiometry, and composition of SVs. The kiss-and-run hypothesis, which argues that vesicles fuse partially with the PM during exocytosis, provides one explanation of how vesicles fully maintain their identity (Ceccarelli et al., 1973). On the other end of the spectrum, vesicular proteins have been shown to diffuse on the PM and re-cluster on sites of endocytosis via the endocytic machinery. Between these two opposing views, our results demonstrate that PM-stranded proteins form nanoclusters via different mechanisms and play a critical role in controlling reuptake of vesicular proteins into SVs.

## Resource availability

### Lead contact

Information and requests for reagents, materials and resources should be directed to and will be fulfilled by Professor Frederic Meunier.

### Materials availability

Syt1-pH was a gift from Volker Haucke. Syt1^K326A/K328A^-pH, SV2A-pH, SV2A^T84A^-pH, SV2A-mCer, SV2A^T84A^-mCer and pSUPER vectors that co-express SV2A shRNA (shRNA sequence - GAATTGGCTCAGCAGTATG) with either mCer or Syt1-pHluorin were described previously (Zhang et al., 2015). SV2A^Y46A^-pH and Syt1-HA were generated as reported in (Harper et al., 2020). mEos4b-Clathrin-15 was a gift from Michael Davidson (Addgene plasmid # 57506; http://n2t.net/addgene:57506; RRID:Addgene 57506). This study did not generate new reagents or plasmids, except for SV2A^Y46A^-mCer and SV2A^T84A/Y46A^-mCer which were generated from either SV2A-mCer or SV2A^T84A^-mCer using the primers forward - GCATCCAGTGATGCT<u>G</u>CTGAGGGCCATGACGAG; Y46A reverse – CTCGTCATGGCCCTCAG<u>C</u>AGCATCACTGGATGC (mutated bases underlined).

### Data and code availability

No original codes are used in this study. A custom-built python tool for NASTIC analysis was can be found at (Wallis et al., 2021) with the Python code available at https://github.com/tristanwallis/smlm_clustering. Analysis of pH fluorescence in nerve terminals was carried out using a custom-made script based on background thresholding. This was used to select nerve terminals, which placed regions of interest of identical size over those responding to stimulation.

## Experimental model and subject details

Experiments were performed using neurons derived from wild-type C57BL/6J mice. For experiments in Brisbane, all work was carried out in accordance with the Australian Code and Practice for the Care and use of Animals for Scientific Purposes and approved by the university of Queensland Animal Ethics Committee (QBI/254/16.NHMRC). All C57/B6 mice were housed with 12 hr light/dark cycle (light exposure between 7am-7pm). Breeders were fed with autoclaved mouse and rat cubes (Specialty Feeds). Adult mice were culled by cervical dislocation. Embryos were euthanised by decapitation. For experiments in Edinburgh, animal work was performed in accordance with the UK Animal (Scientific Procedures) Act 1986, under Project and Personal Licence authority and was approved by the Animal Welfare and Ethical Review Body at the University of Edinburgh (Home Office project licence – 70/8878). All animals were killed by schedule 1 procedures in accordance with UK Home Office Guidelines; adults were killed by cervical dislocation followed by decapitation, whereas embryos were killed by decapitation followed by destruction of the brain. Wild-type C57BL/6J mice were sourced from an in-house colony at the University of Edinburgh. All mouse colonies were housed in standard open top caging on a 14-hour light / dark cycle (light 07:00–21:00). Breeders were fed RM1 chow, whereas stock mice were maintained on RM3 chow.

## Method details

### Hippocampal cell culture

Prior to dissection, 29 mm glass-bottom dishes (Cellvis, CA, USA) were coated in poly-L-lysine (PLL) and left to incubate (37°C, 24 hrs). Hippocampal neurons were dissected from E16 embryos from C57/BL6J mice as previously described (Joensuu et al., 2017). Dissection was carried out in Hank’s buffered salt solution (1X), 10 mM HEPES pH 7.3, 100 U/ml penicillin-100 μg/ml streptomycin. Digestion of hippocampal tissue was carried out using trypsin (0.25% for 10 min) and subsequently halted using fetal bovine serum (FBS, 5%) with DNase I. Suspension was triturated and centrifuged (1500 rpm, 7 min), resuspended in plating medium (100 U/ml penicillin-100 μg/ml streptomycin, 1x GlutaMax supplement, 1x B27 and 5% FBS in neurobasal media). Seeding of neurons was carried out in glass-bottom dishes (1 x 10^5^ neurons covering the central glass bottom of each dish). Subsequently, plating media was fully replaced (2-4 hrs post-seeding) with culturing media (100 U/ml penicillin-100 μg/ml streptomycin, 1x GlutaMAX supplement, 1x B27 in neurobasal media). Hippocampal neurons (DIV13-15) were transfected with plasmid (2 hrs, 2-3 μg per dish) with Lipofectamine2000 (minimum 24 hrs). For pH imaging experiments, dissociated primary hippocampal-enriched neuronal cultures were prepared from E16.5-18.5 embryos from wild-type C57/BL6J mice of both sexes as outlined (Harper et al., 2017; Zhang et al., 2015). In brief, isolated hippocampi were digested in 10 U/mL papain in Dulbecco’s PBS, washed in Minimal Essential Medium (MEM) supplemented with 10 % foetal bovine serum, and triturated to single cell suspension. This cell suspension was plated at 3 - 5 x 10^4^ cells on poly-D-lysine and laminin-coated 25 mm coverslips. Cells were transfected on 7 - 9 days-in-vitro (DIV) with Lipofectamine 2000 as per manufacturer’s instructions (Gordon and Cousin, 2013).

### Super-resolution microscopy

For live single particle tracking, neurons were placed in low K^+^ imaging buffer (5.6 mM KCl, 2.2 mM CaCl_2_, 145 mM NaCl, 5.6 mM D-Glucose, 0.5 mM ascorbic acid, 0.1% BSA, 15 mM Hepes, pH 7.4) at 37°C on a Roper Scientific Ring-TIRF microscope with a CPI Apo 100x/1.49N.A. oil-immersion objective (Nikon Instruments, NY, USA) with a Perfect Focus System (Nikon Instruments). Imaging was carried out using Evolve 512 Delta EMCCD cameras (Photometrics, AZ, USA), an iLas2 double laser illuminator (Roper Scientific, FL, USA), a quadruple beam splitter (ZT405/488/561/647rpc; Chroma Technology, VT, USA) and a QUAD emission filter (ZET405/488/561/640m; Chroma Technology). For imaging molecules on the plasma membrane, Universal Point Accumulation Imaging in Nanoscale Topography (uPAINT) was carried out (Giannone et al., 2010). Neurons were stimulated with high K^+^ buffer (56 mM KCl, 0.5 mM MgCl_2_, 2.2 mM CaCl_2_, 95 mM NaCl, 5.6 mM D-Glucose, 0.5 mM ascorbic acid, 0.1% BSA, 15 mM Hepes, pH 7.4) containing atto647N-labelled anti-GFP nanobodies (NB) (Synaptic Systems) at 3.19 pg μl^-1^. For visualization of Syt1 nanoclusters with clathrin, uPAINT imaging of Syt1-pH-atto647N-NB was carried out in tandem with single particle tracking Photoactivated Localization Microscopy (sptPALM) of clathrin-mEos4b. The mEos4b fluorophore was excited through application of 405 nm laser, which triggered its photoconversion from green to red. Photoconverted mEos4b was simultaneously imaged using excitation with a 561 nm laser.

For imaging of Syt1-pH-atto647N-NB internalised in recycling vesicles, sub-diffractional tracking of Internalised Molecules (sdTIM) was used. Neurons were pulsed for five minutes in high K^+^ buffer containing anti-GFP atto647N-NB (3.19 pg μl^-1^), before being washed with low K^+^ imaging buffer (5x) and left under resting conditions for a further five minutes. Subsequently, neurons were incubated in imaging buffer containing AcTEV protease (Invitrogen, 12575-015) (15 min). For 3D-structured illumination microscopy (SIM), neurons underwent sdTIM imaging and subsequently fixed in paraformaldehyde (PFA) (4% in phosphate-buffered saline/PBS) (15 min), washed in PBS (5x) and mounted in non-hardening antifade mounting medium (Vectashield, H-1000). SIM acquisitions were taken on an Elyra PS.1 microscope (100x objective). A series of z-stacks (23 slices, 0.1 μm intervals, 1024×1024 pixels) were taken sequentially in three channels (488, 561 and 640 nm). For channel alignment (xyz), z-stacks were taken of multifluorescent Tetraspeck beads (Invitrogen, T7279).

### Immunocytochemistry

For immunolabelling of endogenous SV2A, transfected neurons were fixed in PFA (4%) in PBS (20 min). Neurons were washed in PBS (3x) and incubated in blocking buffer of bovine serum albumin (BSA) (1% in PBS) for 30 minutes. Neurons were subsequently incubated with with a rabbit anti-SV2A (ab32942) (1:200, 1 hour) in blocking buffer. Neurons underwent further washes in PBS (3x) and were labelled with anti-rabbit alexa594 (1:1000, 30 minutes), prior to undergoing additional PBS wash steps (3x). Fixation, blocking, antibody incubation and wash steps were all carried out at room temperature.

### Fluorescence imaging

Timelapse recordings of pH were performed as previously described (Harper et al., 2020). Hippocampal neurons were transfected with pH and mCer-tagged fluorescent proteins at DIV7. Between DIV 13-15, neurons were mounted in a Warner Instruments (Hamden, CT, USA) imaging chamber on a Zeiss Axio Observer D1 or Z1/7 inverted epifluorescence microscope (Cambridge, UK) with a Zeiss EC Plan Neofluar 40x/1.30 oil immersion objective fitted with an AxioCam 506 mono camera (Zeiss). The pH fluorophore was excited at 500 nm. SV2A-mCer was excited at 430 nm. Visualisation of mCerulean and pH was carried out using a long-pass emission filter (>520 nm). Field stimulation was carried out on neuronal cultures using a train of 300 action potentials at 10 Hz (100 Ma, 1 ms pulse width). Images of pH were taken at 4s intervals. After 180s post-stimulation, neurons were perfused with an alkaline buffer containing NH_4_Cl (50 Mm), increasing the intracellular Ph thereby revealing the total fluorescence levels of pH.

### Co-immunoprecipitations and western blotting

HEK293 cells were maintained at 37°C, 5% CO_2_ in culture media (MEM (Invitrogen, 41966-029), 10% FBS, 1% penicillin/streptomycin). Cells were co-transfected with SV2A-mCer (WT, T84A or Y46A) and Syt1-HA with lipofectamine2000 (48 hrs). Cells were solubilised in HEPES buffer (50 mM HEPES pH 7.5, 0.5% Triton X-100, 150 mM NaCl, 1 mM EDTA, 1 mM PMSF, protease inhibitor cocktail for 1 hour) prior to centrifugation (17000g for 10 min), from which the resulting supernatant was isolated and treated with GFP TRAP beads (Chromotech, Germany) and rotated at 4°C for 2 hrs, followed by additional wash steps in HEPES buffer (3x). Samples were incubated in SDS sample buffer (10 minutes at 65°C) and loaded on an SDS-PAGE gel for western blotting, which was carried out in accordance with previous studies (Anggono et al., 2006). Primary antibodies used were anti-GFP rabbit (ab6556, 1:4000) and anti-HA rabbit (ICLlab, RHGT-45A-Z, 1:20 000) or rabbit anti-synaptotagmin-1 (Synaptic Systems, #105103, 1:5000). IRDye secondary antibodies (800CW anti-rabbit IgG (#925-32213, 1:10 000); 680RD anti-rabbit IgG (#926-68071, 1:10 000)) and Odyssey blocking PBS buffer (# 92740000) were from LI-COR Biosciences (Lincoln, Nebraska, USA). Blots were visualised using a LiCOR Odyssey fluorescent imaging system (LiCOR Biotechnology, Cambridge, UK). Band densities were determined using LiCOR Image Studio Lite software (version 5.2). The amount of synaptotagmin-1-HA coimmunoprecipitated was normalised to the amount of input protein. These values were then normalised to the amount of immunoprecipitated SV2A-mCerulean.

### Image processing

For single particle tracking, image processing was carried out in PALMTracer, a custom-written software that operates in MetaMorph (Molecular Devices, CA, USA) (Kechkar et al., 2013). Regions of interests (ROIs) were drawn around nerve terminals defined as hotspots of increased pHluorin fluorescence. A spatial resolution of 0.106 μm was set as the detection limit. Trajectories lasting a minimum of eight frames were selected and reconstructed. The MSD was calculated by fitting the equation MSD(*t*)=*a*+4*Dt* (where *D* diffusion coefficient, *a* is y intercept and *t* is time), with MSD quantified over a 200 ms period. The diffusion coefficient was calculated were divided into mobile and immobile populations with a diffusion of log10 > −1.45 μm^2^ s^−1^ considered as mobile (Constals et al., 2015). A custom-built python tool was used to perform Nanoscale Spatiotemporal Indexing Clustering (NASTIC) on our track files to determine the size, density, and apparent lifetime of Syt1 and SV2A nanoclusters. NASTIC generates a series of overlapping, spatiotemporal bounding boxes around trajectories used to determine cluster formation (Wallis et al., 2021). Nanoclusters were thresholded at a radius of 0.15 μm, with anything greater excluded from analysis.

Offline data processing of pHluorin-transfected neurons was performed using Fiji is just ImageJ (Fiji) software (Schindelin et al., 2012). A script based on background thresholding was used to select nerve terminals, which placed regions of interest of identical size over those responding to stimulation (see section on data availability). Average fluorescent intensity was measured over time using the Time Series Analyzer plugin before screening regions of interest using a customised Java program that allows for visualisation of the fluorescent responses and removal of aberrant traces from the data. Subsequent data analyses were performed using Microsoft Excel, Matlab (Cambridge, UK) and GraphPad Prism 6.0 (La Jolla, CA, USA) software. The change in activity-dependent pHluorin fluorescence was calculated as F/F_0_ and normalised to the peak of stimulation.

SIM processing and channel alignment were performed using Zen 2012 SP2 Black (version 11.0, ZEISS). Clusters of Rab5-mRFP were observed along the axons of each neuron. Colocalization of points and surfaces was carried out in IMARIS (version 9.6.0). A series of 3-dimensional surfaces were defined for Rab5 cluster points (561 nm) in each z-stack based on the signal intensity across the neuron and confirmed the cluster points based on the presence of synaptotagmin-1-pHluorin (488 nm). For each surface, the points corresponding to internalised Syt1-atto647-NB molecules (642 nm) were identified.

## Quantitative and statistical analysis

A Students *t*-test was performed for comparison between two groups (Fig. 1–2, Fig. 3A, Fig 3C iv-vi, D i-iv, Fig. 4Ci-iii, Fig.4D iv, Fig. 6–7). A one-way ANOVA was performed followed with a post-hoc Tukey’s test for multiple comparisons for gaussian distribution of residuals (Fig. 3C i-iii, Fig. 4B, Fig. 4D ii, Fig. 5B). For non-parametric analysis assuming no gaussian distribution, Kruskal-Wallis test with Dunn’s test for multiple comparisons was carried out (Fig. 5C-D). The level of significance was set to p <0.05. Error bars represent standard error of the mean (SEM).

## Key resources table

| Reagent/resource | Source | Identifier |
| --- | --- | --- |
| Neurobasal™ Medium | Gibco-Thermo Fisher | 21103-049 |
| B-27™ Supplement (50x), serum free | Gibco-Thermo Fisher | 17504-044 |
| D-(+)-Glucose solution 45% | Merck/Millipore | G8769 |
| Penicillin-Streptomycin (10,000 U/mL) | Invitrogen-Thermo Fisher | 15140-122 |
| Bovogen Australian source Foetal Bovine Serum – Heat Inactivated | Interpath | SFBS-AU |
| Glass-bottom dishes (29 mm) | Cellvis, CA, USA | D29-20-1.5N |
| GlutaMAX™ Supplement | Gibco-Thermo Fisher | 35050-061 |
| Atto647N anti-GFP nanobodies | Synaptic Systems | N0301-AT647N-L |
| AcTEV protease | Invitrogen | 12575-015 |
| Paraformaldehyde, EM Grade 16% | ProSciTech | C004 |
| Dyngo4A | McCluskey et al., 2013 | N/A |
| Rabbit anti-SV2A | Abcam | ab32942 |
| Ascorbic acid | Sigma-Aldrich | A5960 |
| Bovine Serum Albumin (BSA) | Sigma-Aldrich | A8022 |
| Zen Black | Carl Zeiss AG |  |
| Python 2.7 | N/A | https://python.org |
| ImageJ | Schneider et al., 2012 | https://imagej.nih.gov/ij/ |
| Metamorph | Molecular Devices | version 7.10.2 |
| Non-hardening antifade mounting medium | Vectashield | H-1000 |
| Paraformaldehyde, EM Grade 16% | ProSciTech | C004 |
| Rabbit anti-SV2A | Abcam | ab32942 |
| Poly-L-Lysine hydrobromide | Sigma-Aldrich | P2636 |
| Zen Black | Carl Zeiss AG |  |
| Python 2.7 | N/A | https://python.org |
| Rab5-mRFP | Vonderheit & Helenius, 2005 | N/A |
| Clathrin-mEos4b | Addgene | http://n2t.net/addgene:57506 |
| SV2A-shRNA-mCer | Zhang et al., 2015 | N/A |
| SV2A-pHluorin | Zhang et al., 2015 | N/A |
| VAMP2-pHluorin | Miesenböck et al., 1998 | VAMP2-pHluorin |
| Syt1-pHluorin | Zhang et al., 2015 | N/A |

## Author contributions

C.S., C.H., M.A.C. and F.A.M. conceived and designed the project. The manuscript was written by C.S. and F.A.M. Additional manuscript edits were provided by M.A.C., M.J. and T.W. Single particle tracking (uPAINT, sdTIM and sptPALM), 3D-SIM imaging and processing was carried out by C.S. Fluorescence recordings of Syt1-pH and SV2A-pH were performed by C.B.H. and C.K. while E.C.D. performed co-immunoprecipitations. T.W. designed the NASTIC cluster analysis software. IMARIS colocalization was performed by A.M. Technical assistance was provided by R.M.M., N.Y. Drift correction was performed by M.J.

## Acknowledgments

The super-resolution imaging was carried out at the Queensland Brain Institute’s (QBI’s) Advanced Microimaging and Analysis Facility. We thank Dr Adekunle Bademosi, Dr Rumelo Amor, Dr Arnaud Gaudin for technical assistance with SIM imaging and analysis. Many thanks to Dr Nick Valmas for providing the illustrations of Syt1-pH (Fig. 1) and SV2A-pH (Fig. 4), and Dr Alex McCann for assistance with figure 8 and for editorial revisions. Thanks to Ms Barbara Duda for editorial revisions. We thank Clem Jones Centre for Ageing Dementia Research (CJCADR) for its support. Thanks to Professor Phil Robinson for providing the Dyngo4A used in this study. This work was also supported by an Australian Research Council Discovery Project grant (170100125), an Australian Research Council Linkage Infrastructure, Equipment, and Facilities grant (LE130100078), a National Health and Medical Research Council (NHMRC) grant (1139316) to F.A.M. A grant from the Biotechnology and Biological Sciences Research Council (BB/L019329/1) to M.A.C. Research was funded in part by the Wellcome Trust [Investigator Award to MAC (204954/Z/16/Z)]. For the purpose of open access, the author has applied a CC-BY public copyright license to any author accepted manuscript version arising from this submission. M.J. is supported by Australian Research Council Discovery Early Career Researcher Award (DE190100565). R.M.M. was supported by the Clem and Jones Foundation, the State Government of Queensland, and the NHMRC Boosting Dementia Research Initiative. C.S. was supported by the Research Training Program (RTP) Scholarship and a QBI top-up scholarship. F.A.M. is a NHMRC Senior Research Fellow (1155794). We thank Dr. Rona Wilson and Mr. Hamish Runicman for assistance with neuronal culture preparation and Dr. Donal Stewart for the development of a Java program to visualise and screen data. The authors of declare no conflict of interest.

